# Structural plasticity of Atg18 oligomers: organization of assembled tubes and scaffolds at the isolation membrane

**DOI:** 10.1101/2022.07.26.501514

**Authors:** Daniel Mann, Simon A. Fromm, Antonio Martinez-Sanchez, Navin Gopaldass, Andreas Mayer, Carsten Sachse

## Abstract

Autophagy-related protein 18 (Atg18) participates in the elongation of early autophagosomal structures in concert with Atg2 and Atg9 complexes. How Atg18 contributes to the structural coordination of Atg2 and Atg9 at the isolation membrane remains to be understood. Here, we determined the cryo-EM structures of Atg18 organized in helical tubes as well as soluble oligomers. The helical assembly is composed of Atg18 tetramers forming a lozenge cylindrical lattice with remarkable structural similarity to the COPII outer coat. When reconstituted with lipid membranes, using subtomogram averaging we determined tilted Atg18 dimer structures bridging two juxtaposed lipid membranes spaced apart by 80 Å. Together with an AlphaFold Atg18-Atg2 model, we propose that Atg18 oligomers form a structural scaffold coordinating the Atg2 membrane bridge. The observed structural plasticity of Atg18’s oligomeric organization and membrane binding provide a molecular framework for the positioning of downstream components of the autophagy machinery.

## Introduction

Macroautophagy (from here on referred to as autophagy) degrades long-lived proteins, macromolecular complexes and organelles in order to recuperate the molecular building blocks for the cell ^1–5^. In this function, autophagy is critical for cellular maintenance and homeostasis in eukaryotes and dysregulation is implicated in neurodegeneration, cancer, inflammation and infection ^6,7^. During autophagy a double-membrane organelle called autophagosome is formed *de novo* ^8^, engulfs cytosolic contents and fuses with the lysosome _9_. In the meantime, more than 40 gene products have been elucidated, which are organized in six major functional protein complexes: Atg1 kinase ^10^, class III phosphatidylinositol 3-kinase (PI-3K) ^11^, Atg9 lipid scramblase ^12^, the Atg2-Atg18 lipid transfer complex ^13^ and two ubiquitin-like conjugation systems ^14^.

In budding yeast *Saccharomyces cerevisiae*, the Atg1 complex initiates the pre-autophagosomal structure (PAS) by phase separation ^15^. Subsequently, small lipid vesicles containing trimeric Atg9 are delivered to the early isolation membrane (IM) ^16^. Initially, the IM was observed in direct contact with membranes of the endoplasmic reticulum (ER) ^17,18^. Moreover, COPII vesicles and its molecular components were also shown to be involved lipid provision to the autophagosome when the IM is in contact with the ER exit site ^19,20^. While the Atg2-Atg18 complex was found to tether the pre-autophagosomal membrane to the ER ^21^ including Atg9 ^22^, at later stages the Atg2-Atg18 complex was observed localized to the expanding tips of the IM ^23,24^. A recent Atg9 cryo-EM structure revealed the trimeric organization and biochemical activity as a lipid scramblase ^25,26^. Atg2 is a very large 1592 aa rod-shaped protein that was shown to function as an intermembrane lipid transfer protein _27–29_ and has homology to a recently resolved VPS13 structure revealing the principal architecture of a lipid slide ^30^.

Closely linked to Atg2 is Atg18 that early on was shown to be essential for the progression of autophagy ^31,32^. Atg18 belongs to the protein family of β-propellers that bind polyphosphoinositides (PROPPIN). The corresponding primary structure contains a series of conserved WD40 repeats forming the 6 or 7 blades of a β-propeller fold. Initially, the X-ray crystal structures of Hsv2, which is a close relative of the PROPPIN family ^33,34^ and the *Pichia angusta* homolog of Atg18 had been determined ^35^. More recently, the structure of *S. cerevisiae* Atg18 was elucidated in the presence of phosphate and citrate, revealing binding sites for phosphatidylinositol-3-phophate (PI3P) and phosphatidylinositol-3,5-bisphophate PI(3,5)P_2_ (PDB-IDs 5LTD, 5LTG) ^36^. Mutations in conserved binding sites for PI3P and PI(3,5)P_2_ (FRRG motif), affect the progression of the Cvt pathway and autophagy as well as vacuolar morphology ^37,38^ supporting the specific membrane adapter role of Atg18. Moreover, Atg18 contains three up to 120 aa loop regions (i.e. 6AB, 6CD, 7AB) that possess high content of predicted structural disorder and emanate from the blades of the β-propeller fold. The 6CD loop (324-406) of Atg18, for instance, participates in membrane binding ^35,39^ and a subsegment was proposed to form an amphipathic helix in the membrane environment ^40^. Moreover, lipid reconstitution experiments of Atg18 with giant unilamellar vesicles were shown to remodel and tubulate membranes thus mediating membrane scission ^40^. Once bound to membranes, Atg18 was found to undergo oligomer formation as observed by cross-linking mass spectrometry ^35^. In addition to P72R73, the 7AB loop (430-460) of Atg18 was shown to bind to the rod-shaped Atg2 molecule ^36^ that is thought to bridge the PI3P-rich IM on one end with a membrane lacking PI3P like the ER at the opposite end^27^.

In addition to Atg18, Atg21 and Hsv2 are structurally related yeast PROPPIN family members involved in membrane trafficking ^41^. It was found that Atg18 is essential for autophagy and the related cytoplasm to vacuole targeting (Cvt) pathway whereas Atg21 solely for Cvt ^13,37^. The specific PI3P binding and downstream Atg recruitment function of Atg18 and Atg21, however, can be compensated by one another ^42^. Hsv2 was shown to be required for microautophagy of the nucleus ^43^ and was the first of the PROPPINs to be structurally resolved ^33,34^. In higher eukaryotes, Atg18 family proteins are also known as WD-repeat protein interacting with phosphoinositides (PIP) (WIPI) with a total of four isoforms occurring in human cell lines. WIPI1, WIPI2 and WIPI4 have been shown to be directly linked to autophagic progression ^44^. Overexpression of WIPI1 led to large and elongated light microscopic punctae in human cell lines ^45^. WIPI2 recruits the ATG12-ATG5-ATG16L1 LC3 conjugation machinery ^46^ and WIPI4 was demonstrated to biochemically interact with ATG2A and assist in the lipid transfer activity ^28^. The Atg18/WIPI family constitute a complementary network of proteins involved in autophagy and autophagy-related cellular processes.

Although various aspects of Atg18 contributions to the autophagy pathway have been investigated, the involvement and structural role of oligomeric species including Atg2 at the IM remain poorly understood. In this study, we applied electron cryo-microscopy (cryo-EM) to solve structures of helically assembled, soluble and membrane-bound Atg18 using high-resolution cryo-EM structure determination as well as subtomogram averaging. Together with structural models the observed structural plasticity indicates that oligomeric Atg18 dimers when bound to lipid membranes may define the spatial coordination of Atg2 with respect to the IM.

## Results

### Atg18 forms higher-order helical assemblies

In order to structurally characterize Atg18 oligomers and their interaction with lipid membranes, we recombinantly overexpressed *Saccharomyces cerevisiae* Atg18 and subsequently purified it under high-salt conditions. Interestingly, once dialyzed into low salt buffer (100 mM KCl), Atg18 formed large supramolecular structures when observed in negatively stained EM samples. Using cryo-EM of vitrified Atg18, we identified helical tubes of 260 Å diameter including a diamond-shape repeat pattern along the helix (**Fig. 1A**). We acquired cryo-EM micrographs of wildtype Atg18 (Atg18-WT) and Atg18-PR72AA, an Atg2 binding mutant ^47^, and subjected them to segmented helical image processing. 2D class averages of the helical tubes revealed individual Atg18 propeller disc-shaped structures including individual blade separation (**Fig. 1B**). The presence of helical layer lines beyond 1/10 Å in the corresponding Fourier transforms indicated an ordered helical repeat of the assembly. The transforms of Atg18-WT and Atg18-PR72AA ^47^ supported the same structural organization of both helical assemblies (**Fig. 1C**), with an improved helical ordering in the Atg18-PR72AA mutant indicated by the higher diffracting layer line at 1/6 Å. The assemblies exhibited a pitch of approximately 100 Å with 5.1 helical units per turn. Using helical parameters of 19.4 Å helical rise and 70.0° rotation, we determined the helical structures to 3.8 Å and 3.3 Å, respectively (**Fig. 1D-E; Suppl. Fig. 1A-C**). Interestingly, Atg18-WT tubes exhibited diameters of 210 Å whereas Atg18-PR72AA tubes are slightly wider at 220 Å (**Suppl. Fig. 1D-E**). In both structures, the basic helical asymmetric unit contained four copies of Atg18 assembled in a diamond outline with an internal channel of 110 Å diameter.

**Figure 1:**
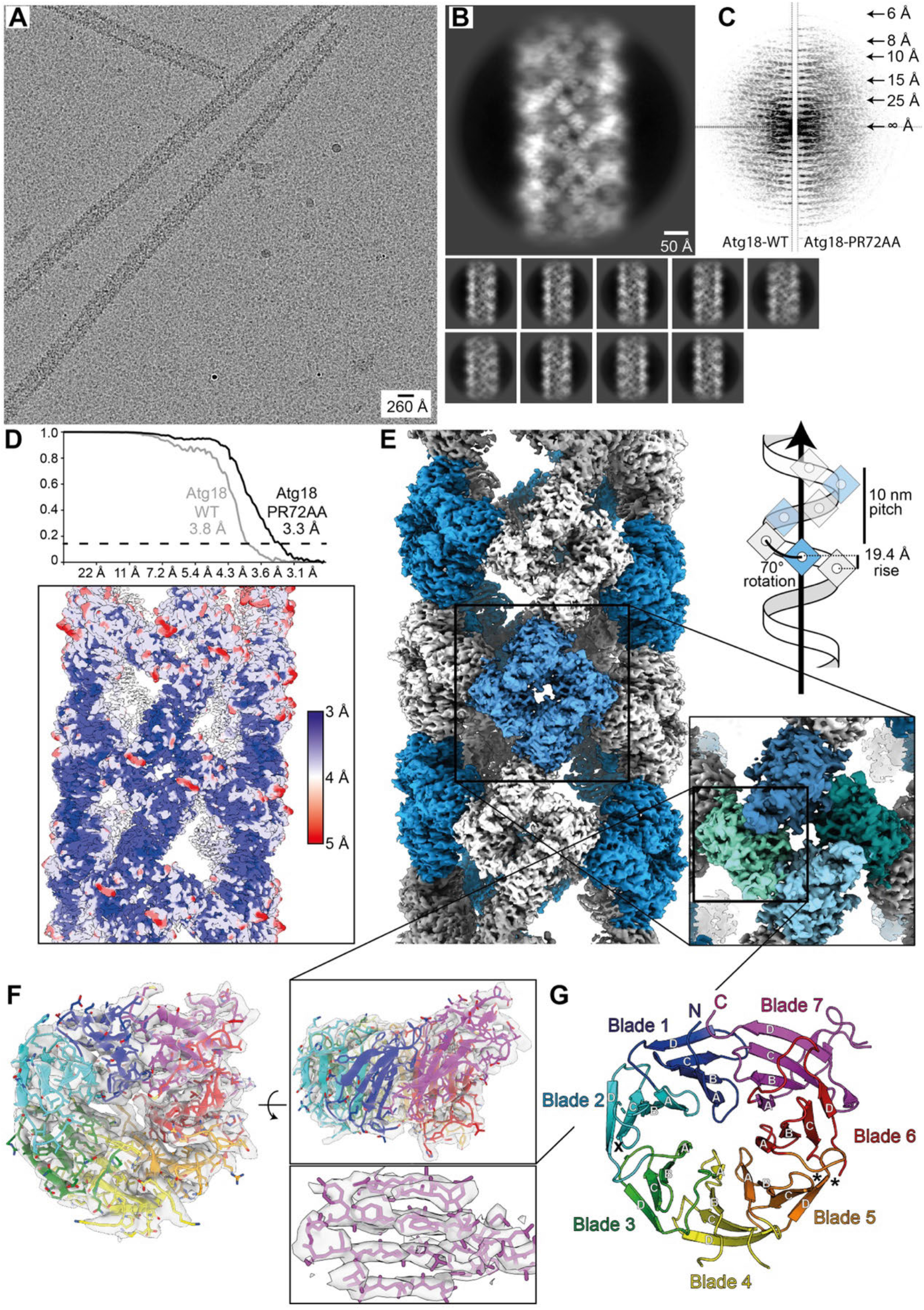
Architecture of helical Atg18 assemblies and cryo-EM structure of Atg18-PR72AA. (A) Micrograph of cryo-vitrified Atg18-PR72AA tubes with a diameter of 260 Å. Note, the grainy background is due to high concentrations of monomeric Atg18. (B) Selected 2D classes and (C) the corresponding Fourier transform including helical layer lines. The panel is composed of the left half of Atg18-WT and right half of Atg18-PR72AA, the latter showing higher resolution diffraction in the power spectrum. (D) Fourier shell correlation threshold at 0.143 indicates global resolutions of 3.8 and 3.3 Å for the 3D structures of Atg18-WT and Atg18-PR72AA, respectively. Local resolution of Atg18-PR72AA mapped onto the 3D density determined by FDR-FSC ^48^ are indicated. (E) 3D density of the helical tube formed by Atg18-PR72AA. The helical symmetry unit is illustrated with alternating blue/white colors (pitch: 100 Å, number of units per turn 5.1; helical rise: 19.4 Å, rotation 70.0°). One diamond-shaped asymmetric unit consists of four Atg18 molecules segmented in blue-green shades in inset. (F) Atomic model refined into the 3.3 Å map (shown as transparent surface) including density β-strand detail of blade 7. (G) Top view of Atg18 ribbon model with the PR72AA mutation located by a cross (x) and the phosphoinositide binding sites in blade 5 indicated by asterisks (*).

When we inspected the cryo-EM maps, they showed the presence of expected density features at the determined resolution such as β-strand separation and side chain densities. The observed features allowed for atomic model building of Atg18 into the cryo-EM density (**Table 1, Fig. 1F-G**). In comparison with the Atg18 crystal structure (PDB-ID 6KYB) ^36^, we found high structural overlap with only minor differences on the rotamer level (RMSD=1.18 Å with 2471 atoms) and slightly extended density at the loop 7AB (**Suppl. Fig. 2A-D**). Residues beyond the β-propeller structural scaffold including the loops 4CD (157-221), 6CD (322-408) and parts of 7AB (446-457) were not resolved in the tube structures of Atg18-WT and Atg18-PR72AA, which is in agreement with both the AlphaFold2 prediction ^49^ (Uniprot ID P43601) and PDB-ID 6KYB. Although the Atg18-PR72AA tube structure was determined at slightly higher resolution when compared with the Atg18-WT (3.3 Å over 3.8 Å), the differences between the atomic models are only detectable as minor positional changes within the assembled tubular structure (**Suppl. Fig. 1C**).

**Table 1.**
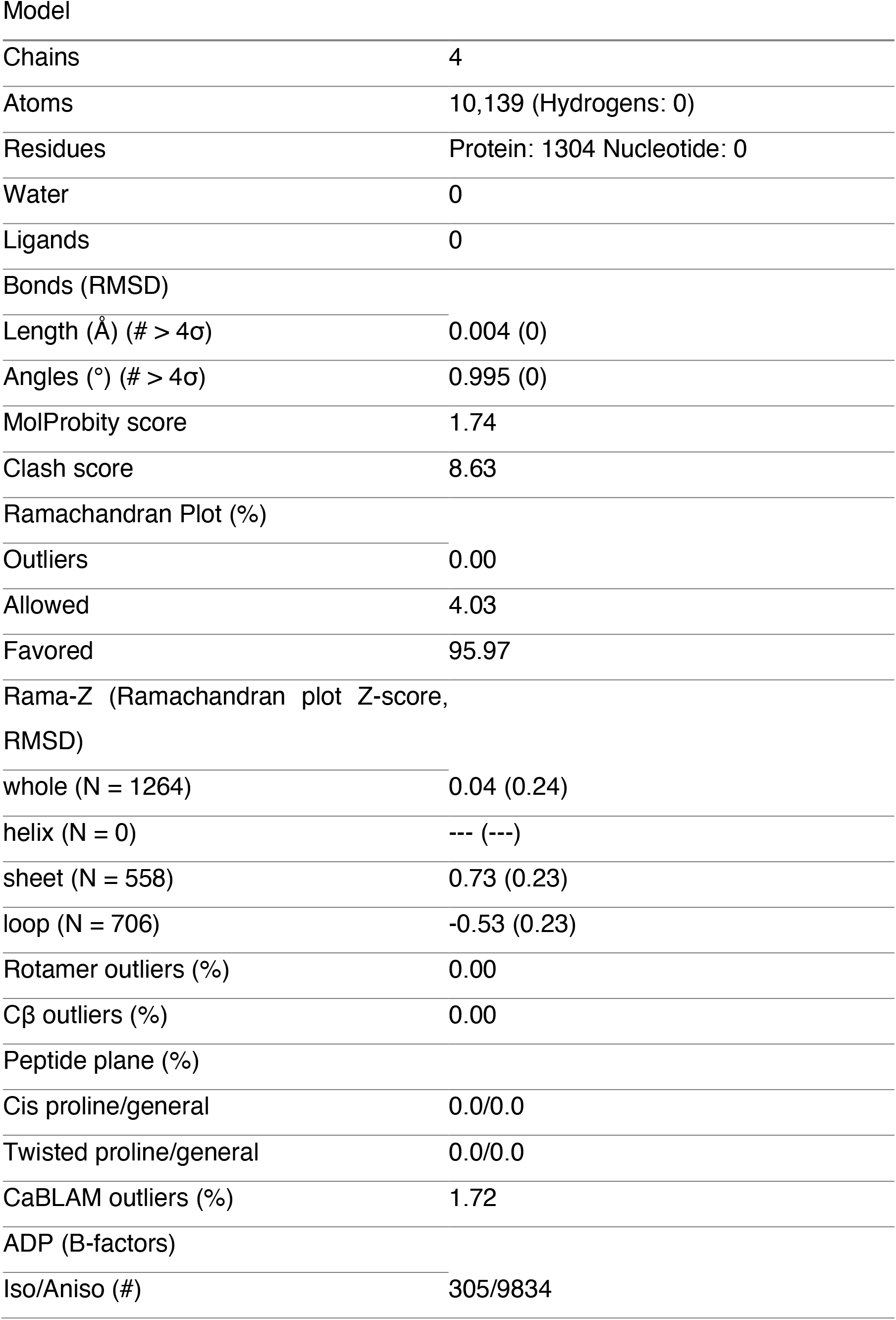
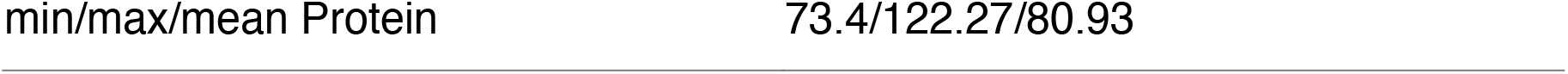
Model report of four Atg18-PR72AA monomers (symmetry unit) built into the 3.3 Å density

|  |  |
| --- | --- |
| Model |  |
| Chains | 4 |
| Atoms | 10,139 (Hydrogens: 0) |
| Residues | Protein: 1304 Nucleotide: 0 |
| Water | 0 |
| Ligands | 0 |
| Bonds (RMSD) |  |
| Length (Å) (# > 4σ) | 0.004 (0) |
| Angles (°) (# > 4σ) | 0.995 (0) |
| MolProbity score | 1.74 |
| Clash score | 8.63 |
| Ramachandran Plot (%) |  |
| Outliers | 0.00 |
| Allowed | 4.03 |
| Favored | 95.97 |
| Rama-Z (Ramachandran plot Z-score, RMSD) |  |
| whole (N = 1264) | 0.04 (0.24) |
| helix (N = 0) | --- (---) |
| sheet (N = 558) | 0.73 (0.23) |
| loop (N = 706) | -0.53 (0.23) |
| Rotamer outliers (%) | 0.00 |
| Cβ outliers (%) | 0.00 |
| Peptide plane (%) |  |
| Cis proline/general | 0.0/0.0 |
| Twisted proline/general | 0.0/0.0 |
| CaBLAM outliers (%) | 1.72 |
| ADP (B-factors) |  |
| Iso/Aniso (#) | 305/9834 |

Common to both structures Atg18-PR72AA and Atg18-WT are two major Atg18 interfaces in the assembly: first, inside the diamond the interface is formed by two perpendicularly arranged Atg18 discs resulting in a “–|” or “T” configuration and, second, the Atg18 interface between diamonds is formed by two Atg18 discs arranged in line giving rise to “– –” or “I” configuration (**Fig. 2A-C**). The T-interface includes R285 and R286 of the FRRG motif that are in direct contact with blade 6 of the adjacent molecule (**Fig. 2D**). The I-interface is located between blade 3 and blade 4, has due to a 180º disc rotation a pseudo dihedral symmetry including in a small tilt of the two discs with respect to each other.

**Figure 2:**
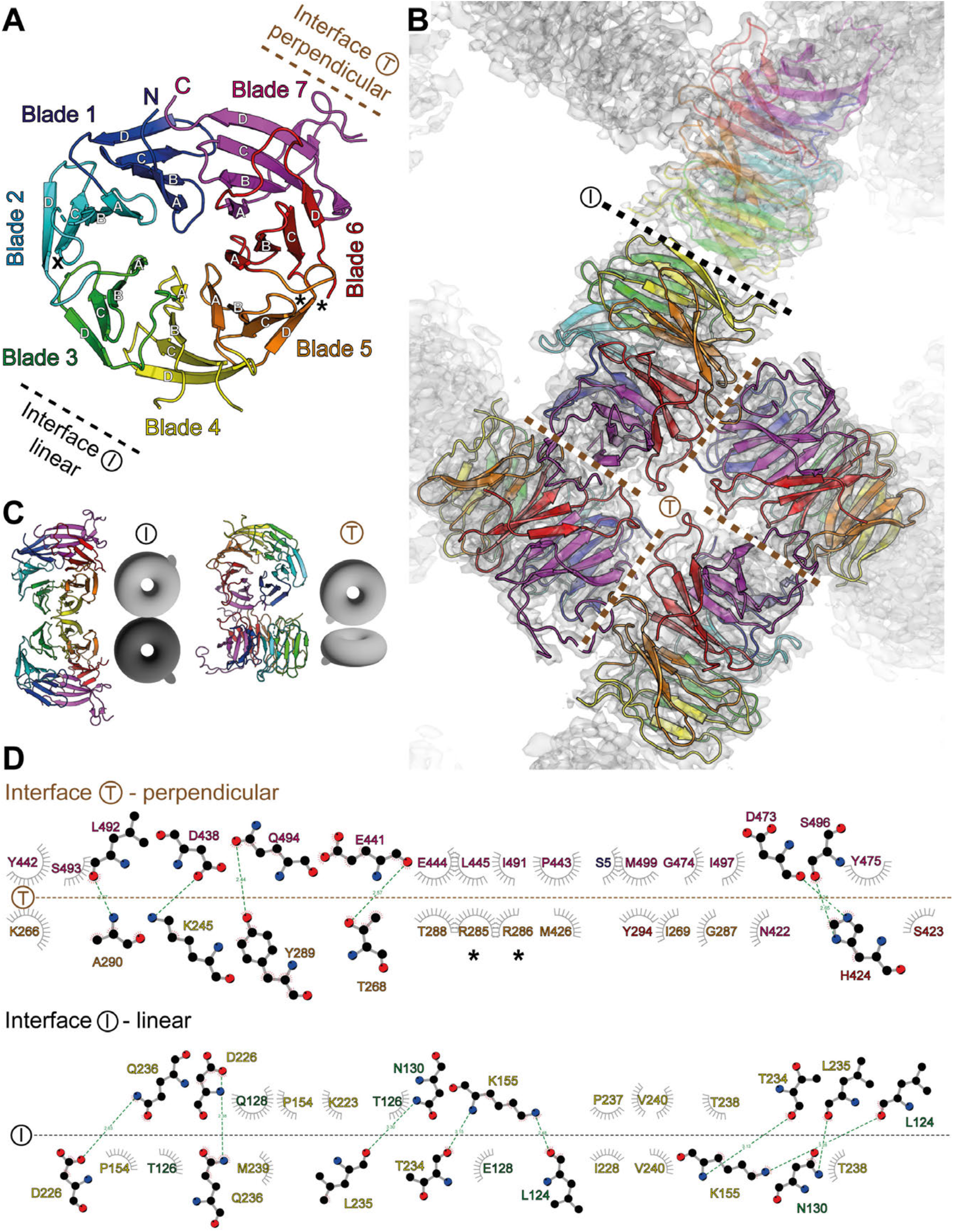
Atg18 assembly interfaces present in the helical tubes. (A) T-Interface (up right/brown) and I-interface (lower left/black) are indicated on the same cartoon color code as in Fig. 1. Location of P72R73 mutation in loop 2BC is indicated by (x). (B) Refined atomic models placed inside 3.3 Å cryo-EM density (transparent background) of Atg18-PR72AA helical assembly. Color code is identical to panel A. T and I-interfaces are indicated with black and brown dashes, respectively. (C) Schematic ribbon presentation of I and T-interface dimers. (D) Schematic residue representations of T and I interfaces. FRRG motif is indicated by asterisks (*).

### Tubular assembly is stabilized by FRRG motif

In order to better understand lipid adaptor function of Atg18 in the context of the newly observed helical assembly, we mapped two binding sites of PI3P and PI(3,5)P_2_ separated by the FRRG-motif onto the presently determined Atg18 assembly structure (**Fig. 3A, B**). Interestingly, the lipid binding sites appear buried in the T-interface of the helical tube suggesting a stabilizing role of the involved residues. In reference to previous mutation studies ^34,37,38^ that abolished binding to PI3P and PI(3,5)P_2_, we replaced the positively charged arginines in the FRRG motif to FGGG by neutral glycines. After purification, we found a complete solubilization of Atg18-FGGG over Atg18-WT tubes when subjected to a pelletation assay (**Fig. 3C** and **Suppl. Fig. 3**). Furthermore, Atg18-FGGG did not form tubes in comparison with Atg18-WT when imaged by negative stain EM. To further experimentally test the influence of the PIP-binding site on the tubular assembly, we added the soluble diC8-PIP derivates of PI3P, PI(3,5)P_2_ and PI(4,5)P_2_ in four-fold molar excess to Atg18-WT prior to low-salt buffer change and visualized these samples by negative stain EM. In contrast to the control-containing formed Atg18 tubes, only smaller particulate aggregates could be observed in the presence of PI3P, PI(3,5)P_2_ and PI(4,5)P_2_, respectively (**Fig. 4D**). Pelletation assays and subsequent separation by SDS-PAGE confirmed that the intensity of Atg18 bands moved to the supernatant fraction for PI(3,5)P_2_ and PI(4,5)P_2_ incubations (**Fig. 4E**). In the case of PI3P, however, we observed a decrease in the pellet fraction in comparison with the untreated control as well as elongated aggregates in negative stain EM. Subsequently, we investigated whether pre-formed tubes were affected by the supplemented lipids. Interestingly, only PI(3,5)P_2_ showed disassembled elongated aggregated tubes when visualized by negative stain EM (**Fig. 4F**), whereas the addition of PI3P or PI(4,5)P_2_ did not affect the helical assemblies visibly. These observations are in line with the highest measured affinities of PI(3,5)P_2_ to the FRRG site ^39^. Together, these experiments show that the FRRG motif of Atg18 takes up a stabilizing role in the formation of the helical Atg18 assembly, which is further supported by binding studies of PI3P derivates that can prevent the assembly of Atg18 tubes and keep Atg18 soluble.

**Figure 3:**
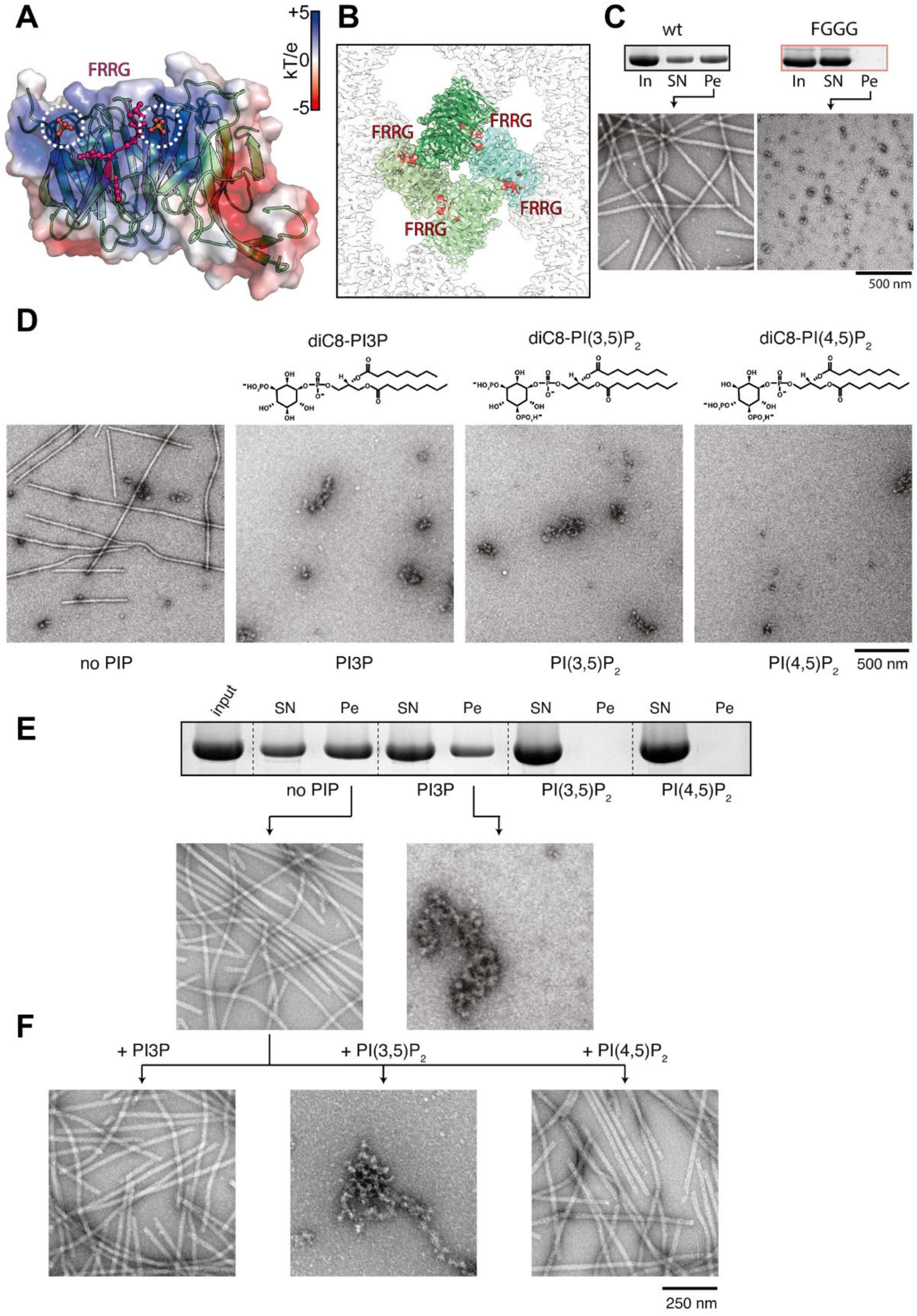
PIP binding by the FRRG motif prevents Atg18 tube formation. (A) Atg18-PR72AA refined atomic model with surface colored by electrostatics, FRRG motif in pink and phosphate binding sites from aligned PDB-ID 5LTD. (B) Diamond shaped tetramer unit (green) with FRRG motifs (red) and their adjacent phosphate binding sites, which are buried in Atg18 tubes. (C) Pelletation assay (top) and negative stain of pellet fractions from Atg18-WT (left) and Atg18-FGGG (right) (D) Top. Chemical formulas of soluble PIP derivates diC8-PI3P, diC8-PI(3,5)P_2_ and diC8-PI(4,5)P_2_ used in binding experiments. Bottom. Negative stain electron micrographs of Atg18 nontreated or treated with 4x molar excess of the indicated PIP derivate preventing tube formation. (E) Pelletation assay with corresponding SDS-PAGE of the supernatant (SN) and the pellet (Pe) fraction and the negative stain EM micrographs of two pellets. (F) Tubes from the negative control were treated with PIP derivates. Only PI(3,5)P_2_ resulted in tube disassembly.

**Figure 4:**
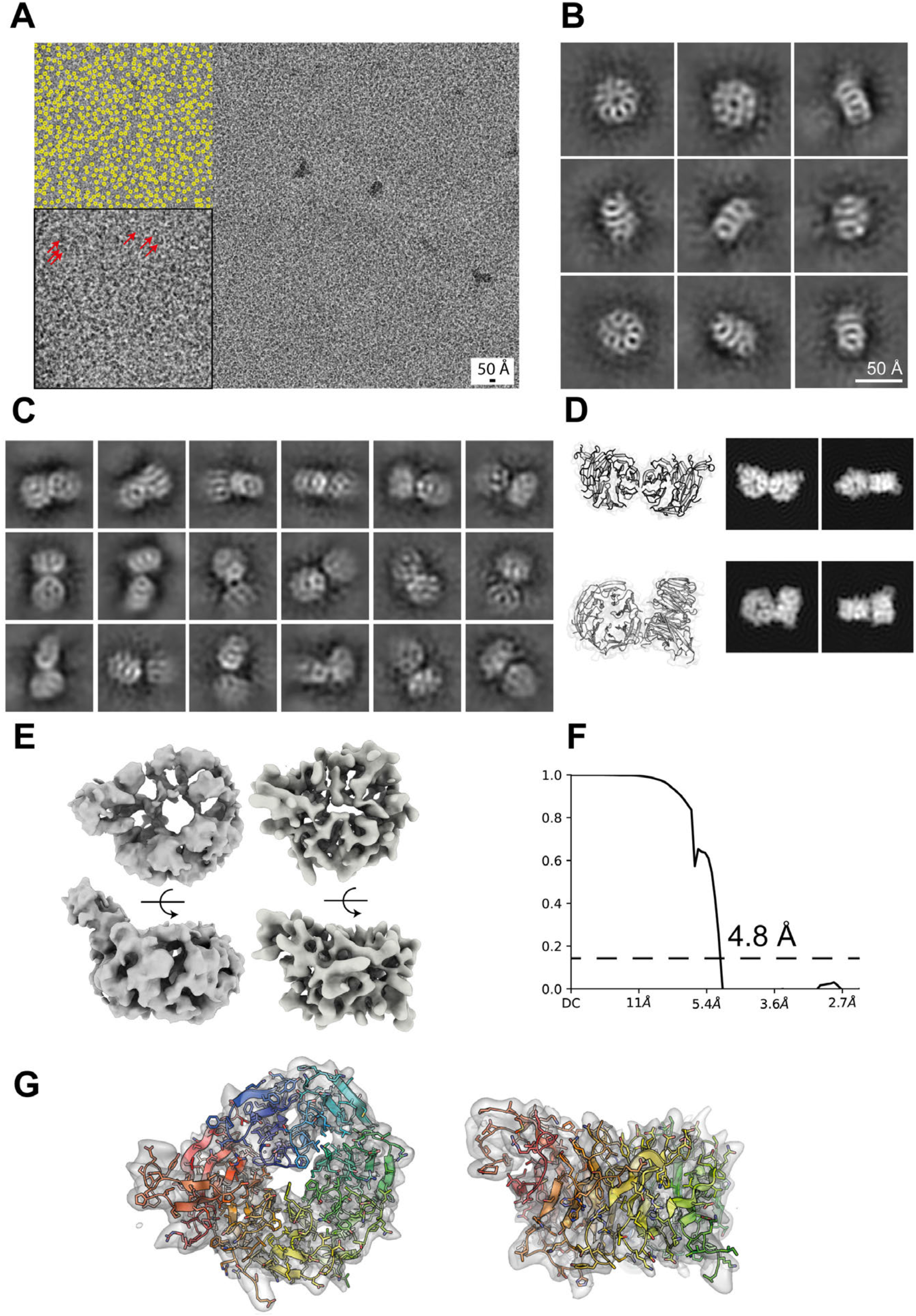
Single-particle cryo-EM structures of soluble Atg18. (A) Typical cryo-micrograph of Atg18-WT with over 1000 particles present (yellow circles in inset). Selected individual molecules are also indicated by red arrows. (B) Representative 2D classes of single Atg18 particle with features of individual blades of the β-propeller. (C) Representative 2D classes of Atg18 dimers from the same dataset. (D) 3D views and 2D projections of I-interface dimer (top) and T-interface dimer (bottom) for comparison with 2D classes in panel C. Some experimental classes are consistent with the packing of I and T-interface dimers observed in the tubular assembly. (E) Ab initio initial 3D model (left) and final 3D structure (right) of Atg18-WT. (F) Fourier shell correlation at the 0.143 threshold indicated global resolution of 4.8 Å. (G) Fitted atomic model taken from Atg18-P72R73 tube structure superimposed with determined 3D structure of monomer Atg18.

### Soluble Atg18 is composed of smaller oligomers

In order to further investigate the structures of soluble Atg18 fractions, we set out to determine structural intermediates of the Atg18 assembly. As indicated by the pelletation assay of Atg18-WT, an equal-share fraction was not pelleted and remained in the supernatant. Using these preparations in high salt buffer, we determined the single-particle cryo-EM structure of soluble Atg18-WT fractions. Due to the very small particle size, i.e. the Atg18 monomer corresponds to 55 kDa, we imaged over 1000 particles on a single cryo-micrograph (**Fig. 4A**). Classification analysis revealed separate views of single Atg18 displaying characteristic structural features of blade separation of the β-propeller (**Fig. 4B**). Alongside Atg18 monomer particles, we were also able to classify a selection of Atg18 dimers (**Fig. 4C**). Comparison with projections of the T and I-interface dimers extracted from atomic model of the Atg18 tube matched some of the 2D classes (**Fig. 4D**). Moreover, we failed to reconstruct coherent dimer 3D structures of these Atg18 dimers presumably due to higher flexibility in solution than observed in the helical assembly. Nevertheless, using monomeric class members we determined the 3D structure of Atg18-WT at 4.8 Å resolution according to the FSC 0.143 criterion (**Fig. 4E**). Next, we successfully docked a single refined atomic model taken from the helically assembled Atg18-P72R73 into the density of monomeric Atg18-WT (**Fig. 4G**). In accordance to the helical assembly structure, three major loop regions were not resolved and disordered in monomeric Atg18. Together, structure determination of soluble Atg18 support the occurrence of monomers and dimers in agreement with the T and I-interfaces observed in the helically assembled lattice structures.

### Visualization of membrane-associated Atg18 oligomers

To further investigate the binding mode of Atg18 to lipid membranes, we mixed soluble Atg18 with PI(3,5)P_2_-doped large unilamellar vesicles (LUVs). Next, we imaged the corresponding plunge-frozen samples by electron cryo-tomography. After acquiring a tilt series with the Volta phase plate, we reconstructed a total of 16 tomograms displaying strong low-resolution contrast. In comparison with common LUV controls that exhibited spherical vesicles of an average 30 nm diameter, the images showed tightly tethered deformed vesicles of square, pentagonal or higher polygonal-like shapes with several 100s nm long stretches of straight and parallel membrane paths next to other vesicles (**Fig. 5A, B**; **Suppl. Fig. 4** and **Suppl. Movie 1**). The membrane surfaces are typically coated by disc-shaped particles corresponding to Atg18 β-propellers. In between parallel membrane stretches, we found density that we interpreted as Atg18. In order to further analyze these membrane-associated structures, we applied membrane segmentation followed by membrane-guided particle picking using the PySeg package ^50^. When generating rotational 2D class averages along the membrane plane from approx. 150,000 extracted subtomograms, some classes showed the presence of additional density between two 60 Å thick bilayers leaving an intra-membrane space distance of 80 Å (**Fig. 5C**). Other classes containing a single membrane only or a second more blurred membrane were excluded from further image processing. Further 3D subclassification resulted in a structure of two S-shaped densities packed against each other made up from eight disc-shaped densities with approx. 50 Å diameter each (**Fig. 5D-F**). A total of 8,300 subtomograms resulted in a structure at 26 Å estimated by mask-less FDR-FSC ^48^ (**Suppl. Fig. 5**). In the density, we found Atg18 molecules in four pairs of dimers with a slightly twisted interface matching the observed I-interface of the tubular Atg18 assembly determined above (**Fig. 5G**). Notably in the cryo-EM density, the basic Atg18 dimer unit was found to assume a 45º tilt angle with respect to both membrane planes, which could be independently confirmed by measurements in raw tomogram slices (**Fig. 5H**). The determined subtomogram structure reveals that tilted Atg18 dimers in I-configuration can establish contact between two opposing bilayers and can further align longer stretches of two opposing lipid membranes in constant distance of approx. 80 Å.

**Figure 5:**
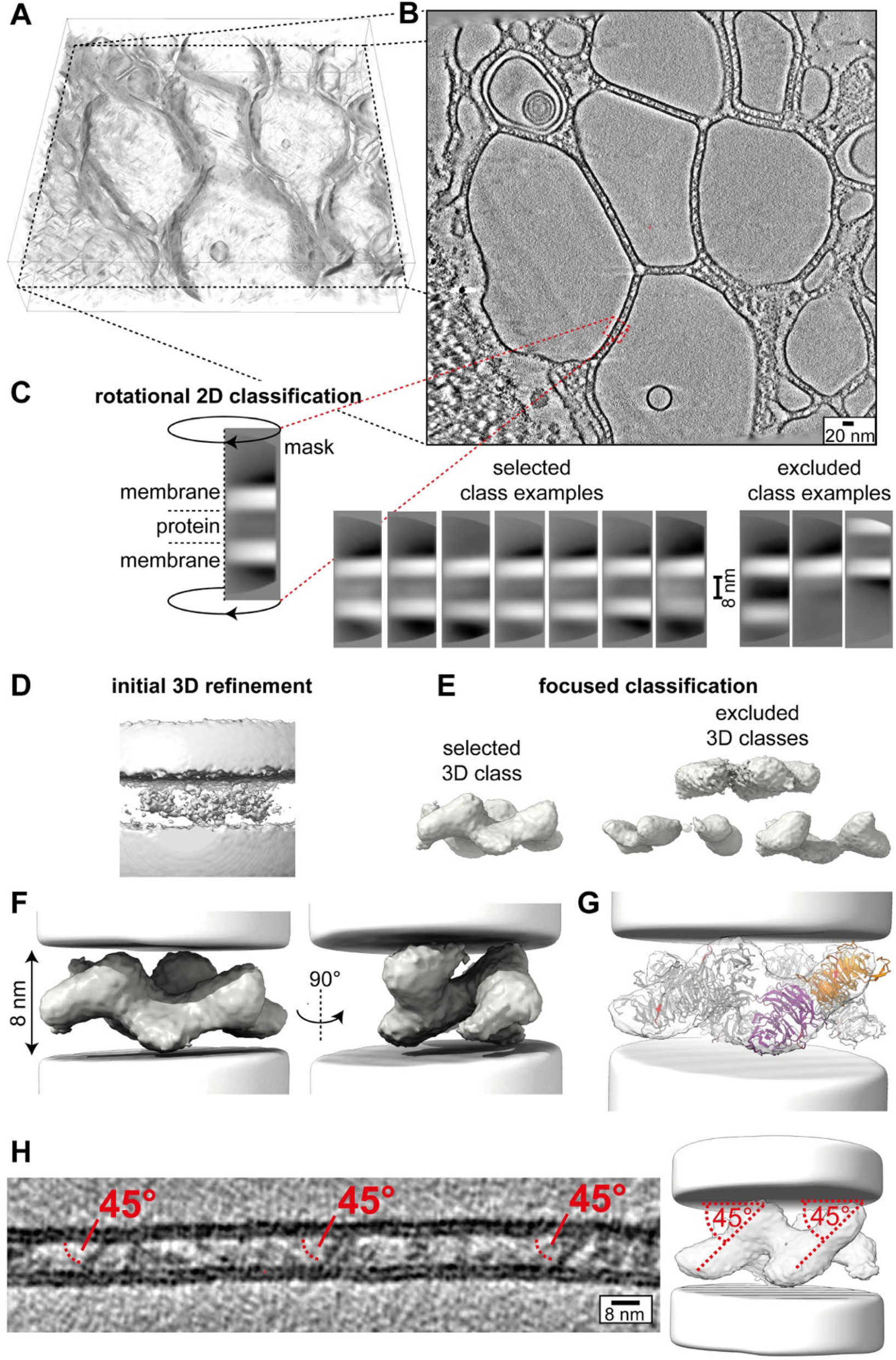
Cryo-ET analysis of membrane-bound Atg18-WT oligomers. (A) Volume density rendering of a reconstructed tomogram containing Atg18-WT and PI(3,5)P_2_-containing large unilamellar vesicles (LUVs). (B) Central slice of the corresponding tomogram acquired with Volta phase plate (averaged five slices to display increased contrast). (C) Examples of rotational 2D classification after subvolume extraction of double-membrane subtomograms. The 60 Å thick membrane bilayers are spaced apart by 80 Å. (D) Initial subvolume average dominated by membrane density contributions. (E) Focused 3D classes after membrane subtraction reveal protein densities. (F) Refined structure of the selected 3D class (C1 symmetry) shows organized stacking of eight disc-shaped molecules. (G) Density fit with four Atg18-WT dimers formed by I-interface previously observed in Atg18 tubular assemblies and soluble dimers. The Atg18 molecule within the dimer is colored orange and magenta, respectively. The remaining six Atg18 molecules are colored grey. FRRG motif is highlighted in red and allows for direct binding of opposing membranes. (H) Slice through reconstructed tomogram reveals Atg18 dimer stacking with the approx. 45° angle relative to the membrane bilayer planes.

### Model of Atg2-Atg18 complex on lipid membranes

In order to further interpret the experimentally obtained subtomogram average structure in the context of the Atg2 complex, we extended the determined Atg18 structures with AlphaFold (AF)-predictions. As a starting point, we verified that the rigid body fit of the Atg18 dimer was consistent with the location of the PI3P binding sites and loop 6CD facing opposite membranes (**Fig. 6A, B**). Based on this structure, we computed the protein complex prediction of N-terminally truncated Atg2 (541-1592) with Atg18 using the AF-multimer approach ^51^ (**Fig. 6C**). The lowest energy structure prediction of the Atg2 (541-1592)-Atg18 complex showed two major interfaces that are consistent with previously determined binding interfaces while leaving the I-dimer interface available for other Atg18 molecules (**Fig. 6D**): first, Atg18-P72R73 had been mutated to disrupt Atg2 binding ^47^ and second, a WIR motif containing peptide of human ATG2A had been crystallized with WIPI3 _52_. This x-Φ-x-Φ-x-x-x-φ-F PROPPIN interaction motif identified in human ATG2A corresponds to residues 921-938 in Atg2 that bind between Atg18’s blade 1 and 2 (**Fig. 6E**). The full-length Atg2 model is available from the AF-EBI databank (Uniprot ID P53855). The Atg2 molecule consists of a 200 Å long β-helix cylinder with a hydrophobic channel inside predicted at high confidence with accessory loops and α-helices predicted at lower confidence. When extending the truncated Atg2 (541-1592) model with the predicted full-length Atg2 and the determined Atg18 dimer, the conserved basic N-terminal residues of Atg2 are in contact with the opposite membrane ^27^ (**Fig. 6F**). In this rigidly extended complex, Atg2 assumes a tilted orientation with respect to the lipid membrane (**Fig. 6 G-H**). Together, based on the experimentally determined Atg18 dimer membrane scaffold, we modeled a Atg2-Atg18 dimer complex in which Atg2 emanates at a tilted angle with respect to the lipid membrane.

**Figure 6:**
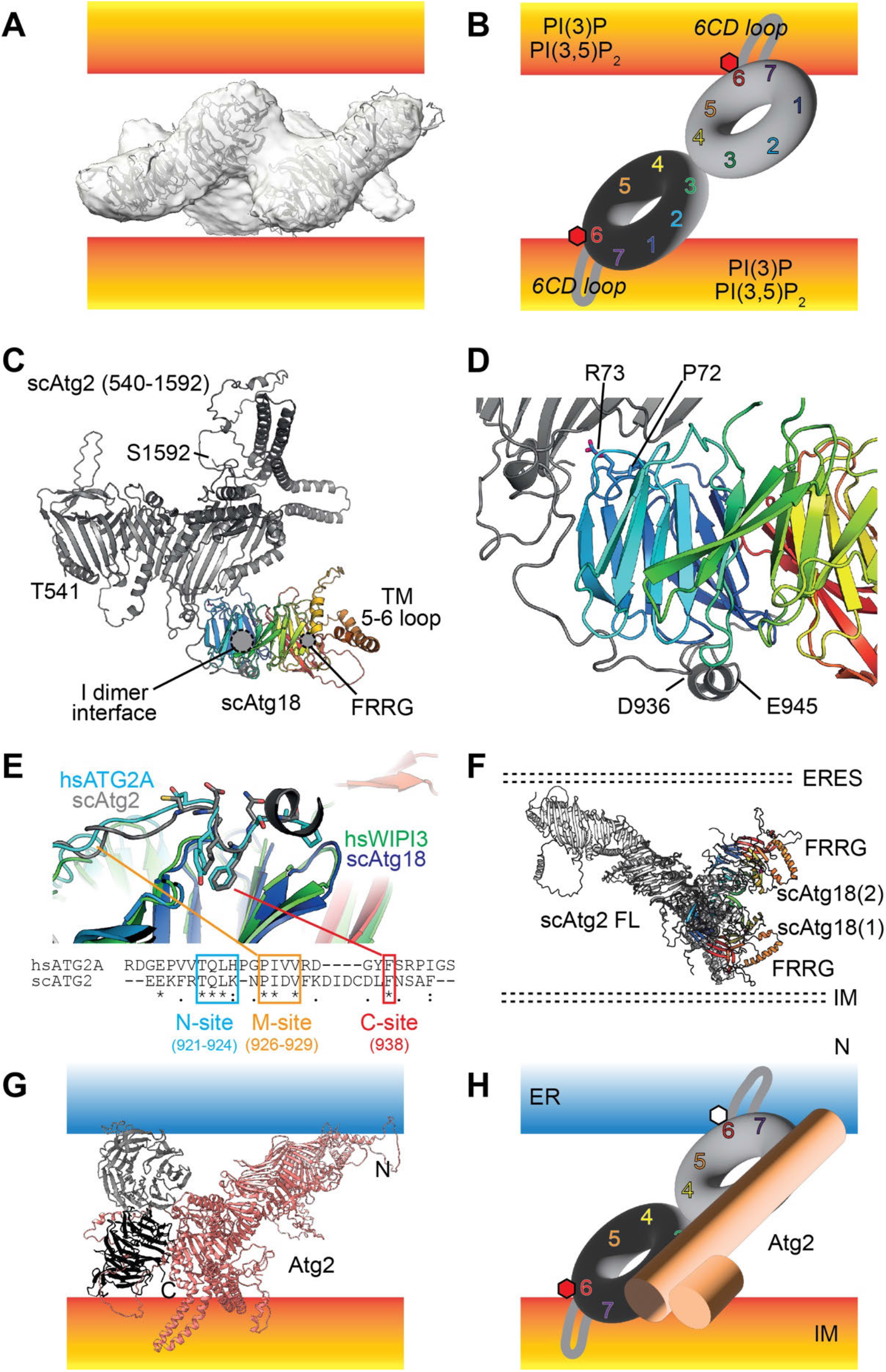
Modeling the Atg18-Atg2 complex on lipid membranes. (A) Cryo-EM subtomogram average density with two fitted Atg18 dimers. (B) Schematic diagram of β-propeller drawn as a ring with assigned propeller blades from 1 to 7. In the fitted dimer loop 6CD and the FRRG motif faces the lipid membrane. The central interface between Atg18 dimer is mediated by loops of blade 3 and 4. (C) Predicted AlphaFold (AF) model of Atg2 (541-1592) with Atg18. (D) Atg2 binds to Atg18 by two interfaces: first, through Atg18’s P72 and R73 ^47^ and second, through blade 1/2 contacting Atg2 at D936 and E945. (E) Structural superposition of ATG2A-WIPI3 (PDB-ID 6KLR) and Atg2-Atg18 model and corresponding sequence alignment shows expected N, M and C-sites of the WIR motif ^52^. (F) When the full-length AF model is placed on the predicted Atg2-Atg18 complex and extended by the observed Atg18 dimer, Atg2 assumes a tilted orientation with respect to the membrane bilayer. (G) Atomic model of Atg18 dimer in complex with Atg2 model derived above including schematic ER (blue) and IM (orange) membranes. Note ER membranes lack PI3P. (H) Consequently, Atg2 when bound to the Atg18 scaffold assumes a tilted orientation with respect to the IM lipid bilayer.

## Discussion

In order to study the structural role of the Atg18 autophagy membrane adapter, we determined the structures of tubular and soluble Atg18 oligomers using high-resolution cryo-EM. Common to these Atg18 structures are two principal oligomeric binding modes that are mediated by so-called T and I-interfaces, respectively. The FRRG motif that is known to bind PIP constitutes a critical part of the T-interface thereby enabling the formation of a large tubular lozenge Atg18 lattice. When Atg18 is reconstituted with lipid membranes containing (PI)3,5P_2_, subtomogram averaging reveals Atg18 oligomers including an elongated tilted Atg18 dimer at the core, which is capable of juxtaposing two opposite membrane bilayers. In the context of a modeled Atg2 complex, the determined membrane-bound Atg18 dimer subtomogram average suggests a scaffolding role of coordinating the structural organization of the Atg2 lipid membrane bridge. The autophagic membrane adapter Atg18 reveals an unexpected structural plasticity in multiple modes of oligomeric organization.

Initial structural characterization of purified Atg18 revealed the formation of higher-order helical assemblies under low-salt conditions (**Fig. 1**). The determined cryo-EM structures of Atg18-WT and Atg18-PR72AA constitute two main propeller interactions mediated by the T and I-interface between neighboring monomers (**Fig. 2**). Analysis of crystal symmetry contacts in a recent X-ray structure revealed identical T and I-interfaces in low-salt buffer, even though the protein was trypsinized to remove the disordered loop regions ^36^. Proteolytically truncated Atg18 forms a tightly packed lattice suitable for 3D crystallization but does not form a hollow cylindrical lozenge lattice built from diamond-shaped tetramers, which we observed for full-length Atg18 in the electron micrographs. Interestingly, the outer membrane-trafficking COPII coat of Sec13-31 displays a very similar structurally related diamond arrangement built from four T-shaped β-propeller units forming a scaffold around lipid membrane tubules ^53^(EMD-11194, PDB-ID 6GZ6) (**Suppl. Fig. 6**). Similar roles of coat formation around lipids can also be envisioned for the observed Atg18 tubes in particular as previous studies showed that Atg18 is capable of tubulating GUVs ^40^. Moreover, recently two groups identified Atg18 independently as a component of the CROP complex in a membrane-associated retromer complex ^54,55^. This architectural conservation of proteinaceous coats suggests functional parallels between different modules of COPII and autophagy trafficking.

The structural similarity of Atg18 tubes to COPII outer membrane coat is a particularly noteworthy observation considering that COPII vesicles have been shown to deliver lipids to the autophagosome IM ^20^. Moreover, phosphorylation states of COPII components affect autophagosome abundance in the cell presumably by modulating coat assembly ^19^. Similarly, when Atg18 was incubated in the presence PIP, we observed the disassembly of the tubular structures into soluble structures suggesting functional precursor role of the resolved tubular assemblies. In fact, the T-interface buries the PI3P/PI(3,5)P_2_ FRRG binding interface and showed direct competition between PIP binding and tube formation (**Fig. 3**). Therefore, our data suggest that the characterized Atg18 tubes act functionally prior to engaging in its adaptor role at the phagophore. In yeast, filamentous assemblies have been shown to accumulate in light microscopic puncta under stress conditions and thereby regulating the activity of metabolic enzymes ^56^. Therefore, a similar storage role may also be envisioned for the observed Atg18 assemblies.

Many other components of the core autophagy machinery such as Atg9 and Atg13 that were shown to be activated and de-activated by chemical modification of phosphorylation ^57,58^. More specifically for Atg18, it was shown for *Pichia pastoris* that Atg18 itself is subject to phospho-regulation by two distinct sites in loops 6CD and 7AB, both reducing PIP-binding affinities ^59^. Under normal conditions in the cell, Atg18 is phosphorylated in loop 6CD and thereby the negative charge prevents the insertion of the amphipathic helix into the membrane ^40^. Similarly, Atg18 phosphorylations in loop 7AB can be expected to affect the stability of the tubular assembly, in particular as the higher resolution structure of Atg2-binding deficient mutant Atg18-PR72AA tubes suggested an increased rigidity over Atg18-WT. Conserved regulatory phosphorylation mechanisms of Atg18 will affect the tube stability and may prime smaller Atg18 oligomers to bind PIP-containing membranes.

Yeast and mammalian autophagosomes exhibit distinct PIP asymmetries, i.e. PI3P is present at the IM while almost absent in the ER membrane ^60^ whereas PI(3,5)P_2_ is found localized in late endosomes, autophagosomes and vacuoles ^38^. The differential binding affinities of different PIP derivatives to Atg18 have been biochemically characterized and quantified before in detail ^39^. In line with the highest affinity, we found that PI(3,5)P_2_ binding leads to disassembly of the helical tubes. The lower binding affinity of PI3P may require additional membrane tethers like the related PROPPIN Atg21 ^42^. Binding to Atg2 through PR72 in loop 7AB ^47^ may contribute further to the destabilization of the tubes in order to assist in the targeted localization of Atg18 to phagophore membranes.

When we investigated soluble Atg18 fractions at high-salt conditions by single-particle cryo-EM, we confirmed the presence of monomers and dimers in line with previous native mass spectrometry characterizations ^35^ (**Fig. 4**). Mass spectrometry cross-linking identified residues in loops 4 BC and loop 6CD engaged in Atg18 oligomer interactions ^35^ consistent with the here determined T and I-interface. Furthermore, the study also revealed that the interaction patterns between Atg18 oligomers changes in the presence and absence of liposomes ^35^. In support, when we incubated Atg18 with PIP-containing membranes, we observed in subtomogram average structures of liposome decorated dimers and tetramers that lack the interaction via the T-interface due to PIP binding mode (**Fig. 5**). Together, Atg18 displays an unexpected structural plasticity of different multimer assemblies depending on buffer condition as well as binding partners, which is mechanistically accomplished by two distinct binding interfaces.

WD40 domains constitute a universal protein interaction scaffold within multiprotein complexes involved in many different biological structures such as histone and ubiquitin binding modules as well as membrane trafficking complexes ^61–64^. In autophagy trafficking Atg18, Atg21 and Hsv2 as well as WIPI1-4 belong to the family of PROPPINs of yeast and mammalia, respectively. Given the ability of Atg18 to form oligomeric structures, it is conceivable that this structural property is conserved in higher eukaryotes. Upon starvation and co-transfection of GFP-WIPI1 in human cells, large cellular punctate structures including lasso filaments were observed by fluorescence microscopy ^45^ compatible with the here observed Atg18 tubes. In higher eukaryotes, ATG2A is known to interact with WIPI1 and WIPI4 whereas WIPI2 contacts ATG16L ^46,65^ another component of the LC3 lipidation machinery. In yeast, Atg18 and Atg21 co-localize to the PAS and Atg18 forms a complex with Atg2 whereas Atg21 contacts Atg16 of the Atg8 lipidation machinery ^42^. Given the complementary functions and overlapping cellular locations of the WIPI proteins, it will require further investigation whether WIPI proteins form hetero-oligomeric complexes in addition to the observed homo-oligomeric Atg18 structures.

In order to put the determined Atg18 dimer structure bound to the lipid membrane in the context of the Atg2 complex, we computed an AF model of the Atg18-Atg2 complex (**Fig. 6**). Here, Atg18 formed two contacts with Atg2 through P72R73 (loop 2BC) and blade 1/2 _47,52_. In an expanded Atg2-Atg18 dimer complex, the I-interface as well as the FRRG motif in blade 5 including loop 6CD are spatially accessible to engage in oligomer and membrane binding. In the Atg18 scaffolding configuration, the rod-shaped 20-nm long Atg2 molecule assumes an angle of approx. 45º with respect to the membrane bilayer bridging membrane distance of ∼10 nm. *In vitro* reconstitutions of Atg2 and ATG2A increased lipid transfer across vesicles in the presence of Atg18 and WIPI1/4, respectively ^27,28^. Therefore, it is tempting to speculate that the derived geometric arrangement is energetically favorable to immerse Atg2’s N-terminal and C-terminal ends of the hydrophobic lipid channel in the outer membrane leaflets. Moreover, participating complex partners such as lipid scramblase Atg9 may also require binding interfaces at this angle in order to channel lipids more efficiently across the membrane bridge to the growing IM.

## Acknowledgments

The authors gratefully acknowledge the computing time granted by the JARA Vergabegremium and provided on the JARA Partition part of the supercomputer JURECA at Forschungszentrum Jülich ^64^. The authors acknowledge Wim Hagen (EMBL Heidelberg) for assisting in setting up the tomography acquisitions of Atg18 liposome mixtures.

## Author Contributions

S.A.F., D.M. and C.S. designed the research. N.G. and A.M. initially purified Atg18 and identified conditions for Atg18 tube formation. S.A.F. purified protein performed biochemical experiments such as pelletation assay and lipid binding studies. S.A.F. and D.M. determined the presented cryo-EM structures of Atg18. D.M. and A.M-S computed the subtomogram average of membrane-associated Atg18. D.M. and C.S. wrote the manuscript with major input from S.A.F and all other authors.

## Data availability statement

The atomic models of tubular Atg18-PR72AA and Atg18-WT were deposited in the Protein Databank (PDB-ID 8AFY and PDB-ID 8AFW), accompanied with the corresponding cryo-EM maps (EMD-15412 and EMD-15410). Single particle maps of monomeric Atg18-WT and subtomogram average are available in the EM Databank linked to fitted PDB coordinates, respectively (EMD-15410, PDB-ID 8AFW).

## Methods

### Protein expression and purification

N-6xHis-tagged Atg18 (Uniprot ID P43601), Atg18-FGGG and Atg18-PR72AA included in pEXP5-NT/TOPO vectors with ampicillin resistance were recombinantly expressed in *Escherichia coli* BL21(DE3) cells. Briefly, after cell growth to OD_600nm_ of 0.6 the temperature was lowered to 20°C and expression was induced by 0.2 mM IPTG overnight. Cells were harvested at 6,000 *g* and resuspended in lysis buffer (50 mM TRIS, 500 mM NaCl, 10 mM KPO_4_, 10 mM imidazole, 1 mM DTT, 0.5 µg/ml DNaseI, 0.1% Triton X-100, pH 7.5). Cells were lysed by homogenization in a fluidizer (EmulsiFlex-C3, Avestin or CF1 cell disrupter, Constant Systems) using 3 passes. Cell debris was cleared by centrifugation at 40,000 *g* for 45 min at 4°C and supernatant was applied to His-tag affinity purification using Ni-NTA agarose (beads or 5 ml column, Qiagen) in buffer A (50 mM TRIS, 500 mM NaCl, 10 mM KPO_4_, 10 mM imidazole, 1 mM DTT, pH 7.5) and eluted in the same buffer supplemented with 250 mM imidazole. The N-6xHis-tag was cleaved with home-prepared TEV protease, imidazol was removed overnight by dialysis and the cleaved 6xHis-tag was removed by a second round of His-tag affinity purification. The flow-through and wash fractions were pooled and concentrated to ∼1 mg/ml using a 30 kDa cutoff spin concentrator (Amicon) before loading to a pre-equilibrated (50 mM TRIS, 500 mM NaCl, 10 mM KPO_4_, 1 mM DTT, pH 7.5) gel filtration column (HiLoad 16/600 Superdex 200 pg, GE Healthcare). Peak fractions containing Atg18 were pooled and concentrated using a 30 kDa cutoff spin concentrator to 8-10 mg/ml. Protein aliquots were snap frozen in liquid nitrogen and stored at -80°C.

### Atg18 tube formation

Atg18 aliquots were thawed and diluted to 100 µM with tube buffer (20 mM HEPES, 100 mM KAc, 1 mM DTT, pH 7.2) and dialyzed against the same buffer overnight (10 kDa membrane, ThermoScientific or Qiagen) at room temperature. Precipitates were cleared by centrifugation at 2,000 *g* for 10 min and the pellet was disposed. To investigate tube formation and pelletation behavior of Atg18 variants or to enrich Atg18 tubes, 100-150 µl of sample were centrifuged for 120 min at 50,000 *g* and 18°C. The supernatant was carefully removed and the pellet rinsed once with 150 µl tube buffer without detaching the pellet from the tube bottom. For semi-quantitative analysis of the pelletation, the pellet was resuspended in the same volume as the input to the pelletation and identical volumes of input, SN and pellet were analyzed by SDS-PAGE. For EM or other experiments, the pellet was resuspended in the desired volume.

### Soluble lipid and liposome binding experiments

To test the effect of soluble lipids on Atg18 tube formation, diC8-PI3P and diC8-PI(3,5)P_2_ (Echelon) were reconstituted in water (1 mM final concentration) and added in 4x molar excess to Atg18 in gel filtration buffer prior to the dialysis into tube buffer. As a control, the same procedure was performed with water instead of reconstituted lipids. To test the effect of soluble lipids on preformed Atg18 tubes, the soluble lipids in water were added to preformed Atg18 tubes in 4x molar excess. As a control, the same procedure was performed with water instead of reconstituted lipids. LUVs were prepared with a lipid content of 95% DOPC and 5% DO-PI(3,5)P_2_ (Avanti Polar Lipids). DOPC was dissolved in chloroform whereas DO-PI(3,5)P_2_ was chloroform:methanol:water (20:9:1 v/v). Lipid films were generated in a glass vial by evaporating the solvent under a gentle stream of argon followed by vacuum desiccation. The lipid films were rehydrated in LUV buffer (20 mM HEPES, 50 mM NaCl, pH 7.1) for 60 min at 37 °C, vortexed for 30-60 s and transferred to an Eppendorf tube at a final concentration of 1 mg/ml. LUVs were formed by three freeze-thaw cycles in liquid nitrogen and a 30 °C water bath. LUVs were snap-frozen in liquid nitrogen and stored at -20 °C until further use.

### Negative stain electron microscopy

300 mesh Gu grids with predeposited continuous carbon were purchased from Electron Microscopy Science or a homemade 5-10nm thick carbon film (Leica EM ACE600) was floated manually onto 400 mesh Cu/Rh grids (Plano GmbH). Grids were glow discharged in air plasma (0.45 mbar) for 45 s (Pelco EasiGlow) before use. A total of 3.5 µl of the respective sample were applied to the carbon side of the grid, adsorbed for 2 min and gently blotted from the side of the grid with filter paper (Whatman 1). Grids were washed twice with drops of sample buffer followed by immediate blotting. Staining of the adsorbed sample was performed with 2% uranyl acetate solution twice, followed by blotting and drying for at least 15 min before imaging. Images were taken on a FEI Morgagni 268 operated at 100 kV equipped with a 1k CCD camera (Soft Imaging Solutions), on a Philips Biotwin CM120 operated at 120 kV equipped with an identical camera or on a FEI Tecnai Spirit operated at 120 kV equipped with a 4k CCD camera (Gatan UltraScan 4000)

### Electron cryo-microscopy

A total of 3 µl 20 µM Atg18 tube solution in tube buffer (20 mM HEPES, 100 mM KAc, 1 mM DTT, pH 7.2) was applied to freshly glow discharged UltrAuFoil R1.2/1.3 300 mesh grids (Quantifoil) and plunge frozen in liquid ethane using a Leica EM GP (18°C, 99% humidity, 1.5 s single blot) or a ThermoFisher Vitrobot Mark IV (18°C, 100% humidity, 1 s single blot, blot force 6). A total of 3.6 µl 18 µM monomeric Atg18 solution in high salt buffer (50 mM TRIS, 500 mM NaCl, 10 mM KPO_4_, 1 mM DTT, pH 7.5) was applied to freshly glow discharged R1.2/1.3 Cu200 grids (Quantifoil), blotted for 7s (blot force -5, 4°C, 95% humidity) and immediately plunge frozen in liquid ethane in a Vitrobot Mark IV (ThermoFisher). LUVs were prepared as described above. For tomography of Atg18 on LUVs, 1 µl Atg18 in 20 mM HEPES, 150 mM NaCl, pH 7.2 was added to 9 µl of LUVs in LUV buffer. The final Atg18 concentration was 3 µM, LUVs at 0.1 mg/ml. A total of 3 µl of the solution was applied to freshly glow discharged R2.2 Cu300 grids (Quantifoil), blotted from the back side for 1.2s at 20°C and 99% humidity and immediately plunge frozen in liquid ethane using a Leica EM GP. 16 tomograms were recorded in focus with Volta Phase Plate induced phase shifts in dose symmetric scheme starting at 0° in +/-3° increments to 60° with 3e^-^/Å²/tilt fluence. Data collection was performed on 300 kV Titan Krios (ThermoFisher) or 200 kV Talos Arctica (ThermoFisher) instruments in SerialEM ^66^ or EPU software (ThermoFisher). Data collection parameters are summarized in **Table 2**.

**Table 2.**
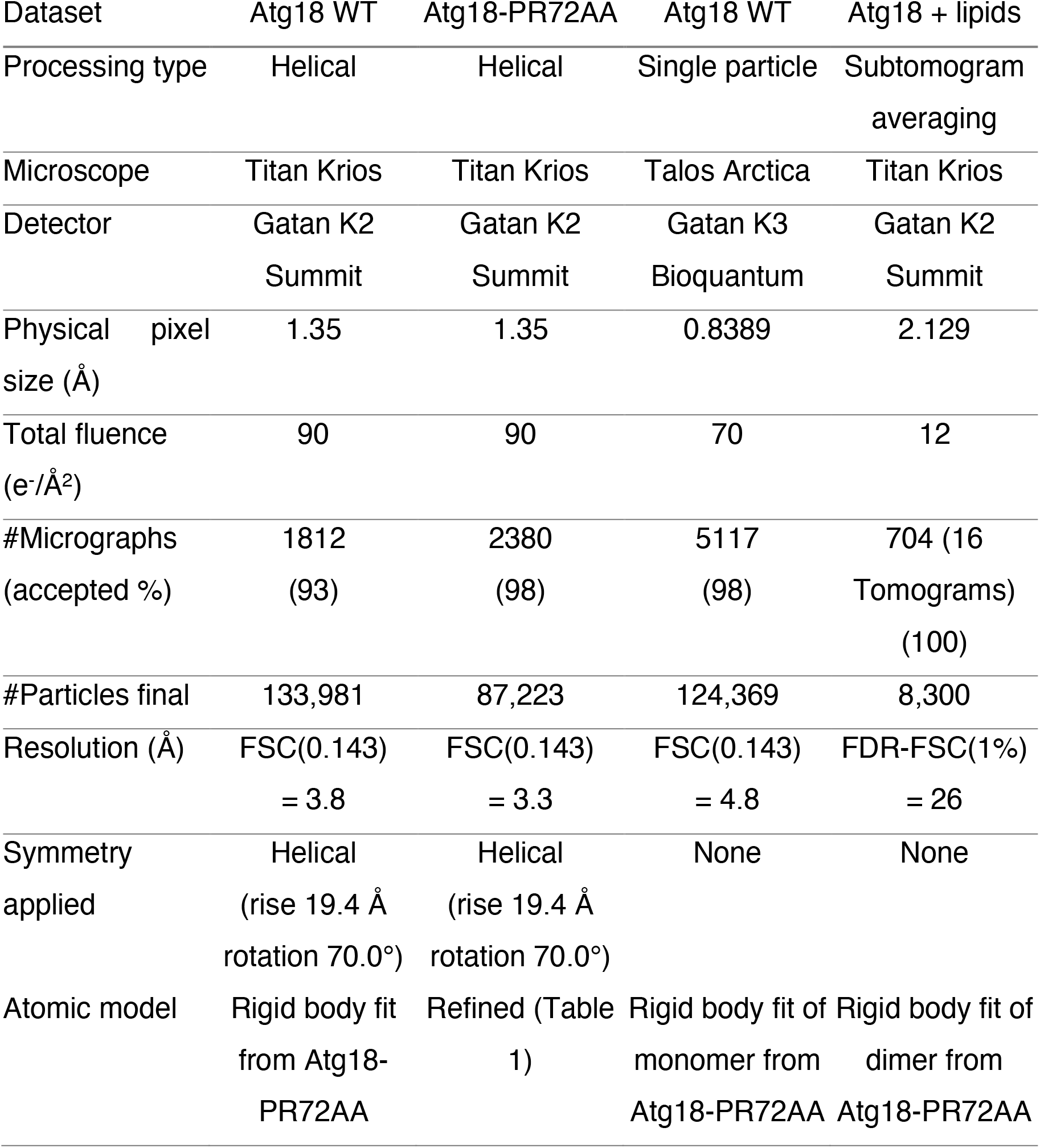
Acquired datasets

### Cryo-EM image processing

Micrographs containing Atg18 tubes and soluble monomer/oligomers were processed in CryoSPARC v3.3.1 ^67–69^. Movies were patch motion corrected, dose weighted and binned to physical pixel size. Patch CTF estimation was performed and micrographs with poor CTF fit resolution >6 Å were excluded (see % used in **Table 2**). Helical tubes were automatically traced (diameter 260 Å, 0.3 (78 Å) step distance), segmented in 400 px / 540 Å boxes and 2D classified. High-resolution 2D classes containing propeller blades were selected. Initially, a symmetry search was performed in the SPRING software package using SEGCLASSRECONSTRUCT ^70^, resulting in symmetry values for pitch and number of units per turn of 99.8 Å and 5.1. The selected classes were also subjected to helical refinement starting with a featureless cylinder without symmetry parameters, resulting in a ∼5 Å 3D reconstruction. Symmetry search in CryoSPARC yielded a global minimum at phi=70°, dz=19.4 Å. Helical symmetry was applied in a second helical refinement run, with on-the-fly refinement of local defocus and global beam aberrations, resulting in 3D reconstructions at FSC(0.143)=3.3 Å for Atg18-PR72AA and FSC(0.143)=3.8 Å for Atg18-WT. 3D classification and 3D variability analysis did not improve the obtained map resolutions. Soluble monomers/oligomer data was picked by template matching against circular blobs (40-60 Å) in CryoSPARC live. Particle extraction was performed in 256 px / 215 Å boxes to include delocalized signal. 2D classification in 200 classes was performed with increased batch sizes (200) and 100 iterations to a high-resolution limit of 4 Å. Monomeric and dimeric classes were split and subjected to *ab initio* 3D map generation with increased resolution ranges (12 Å to 6 Å) and increased batch sizes (300 initial, 1000 final size) with K=3 classes. The monomeric class was low-pass filtered to 8 Å and subjected to 3D refinement and non-uniform local refinement ^68^, resulting in a map with FSC(0.143)=4.8 Å whereas dimeric classes could not be resolved further.

Tomograms were reconstructed using IMOD/eTomo software ^71^ including patch tracking and weighted back projection. Subsequently, they were subjected to the PySeg pipeline ^50^ for particle picking and classification. Briefly, lipid membranes were segmented by TomoSegMemTV software ^72^ and PySeg’s tracing routines were used to pick subvolumes near the segmented membranes. A total of 150,000 picked subvolumes were averaged in three batches and subjected to rotational 2D classification focused by a cylinder mask using the affinity propagation clustering algorithm. Double membrane classes with protein signal in between the membranes were selected and all other classes were excluded (55 classes out of 300 classes included). This way, the majority of the subvolumes were excluded as they contained single membrane classes, double membrane classes with the second membrane outside of the focused mask as well as triple membranes with no connecting densities. 2D classification and class selection was repeated another time with a mask suppressing the membrane signal (32 out of 516 classes included) before extracted subtomograms were subjected to Relion v3.0 software for subtomogram averaging^73^. An *ab initio* 3D model was generated and refined and the PySeg tools were used for masked subtraction of the two membranes (100% dampening). Focused classification into four 3D classes showed one dominating class that was used for 3D refinement. FDR-FSC was used to determine the nominal resolution of 26 Å of the averaged subvolume map ^48^. Atg18 dimers from helical assembly were docked into the map using ChimeraX software^74,75^.

### Atomic model building and refinement

PDB-ID 6KYB was docked into the reconstructed cryo-EM density of Atg18-PR72AA after map auto-sharpening in Phenix v1.20.1 in ChimeraX software. The atomic model of Atg18-PR72AA was manually mutated and slightly modified in Coot v0.9 ^76^ before real-space refinement in Phenix was performed with default parameters^77^. Validation parameters are displayed in **Table 1** ^78,79^. Based on the refined atomic Atg18-PR72R73 model, we used the corresponding Atg18-WT model to rigidly place it into Atg18-WT tubules, Atg18 monomer as well as Atg18 subtomogram average dimer densities.

## Supplementary Figures

**Suppl. Fig. 1:**
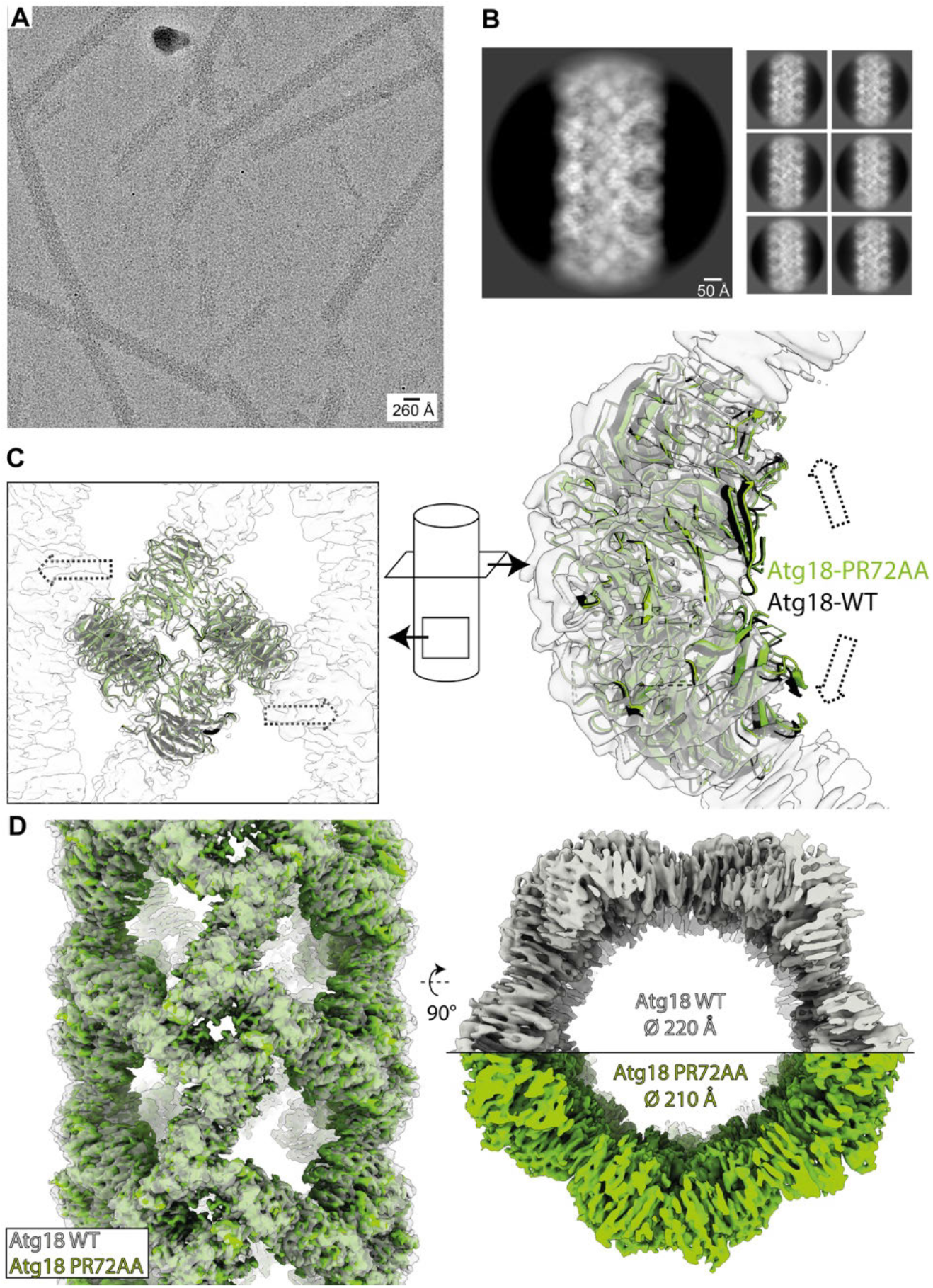
Helical structure of Atg18-WT in comparison with Atg18-PR72AA. (A) Typical electron cryo-micrograph of Atg18-WT tubes. (B) Representative 2D classes of Atg18-WT. (C) Comparison of molecular models of Atg18-WT (black) and Atg18-PR72AA (green) with the Atg18-WT cryo-EM map superimposed in transparent grey. (D) Comparison of respective cryo-EM densities reveals slightly wider diameter of Atg18-WT compared with Atg18-PR72AA. Note that the data were acquired on the same microscope with the same pixel size.

**Suppl. Fig. 2:**
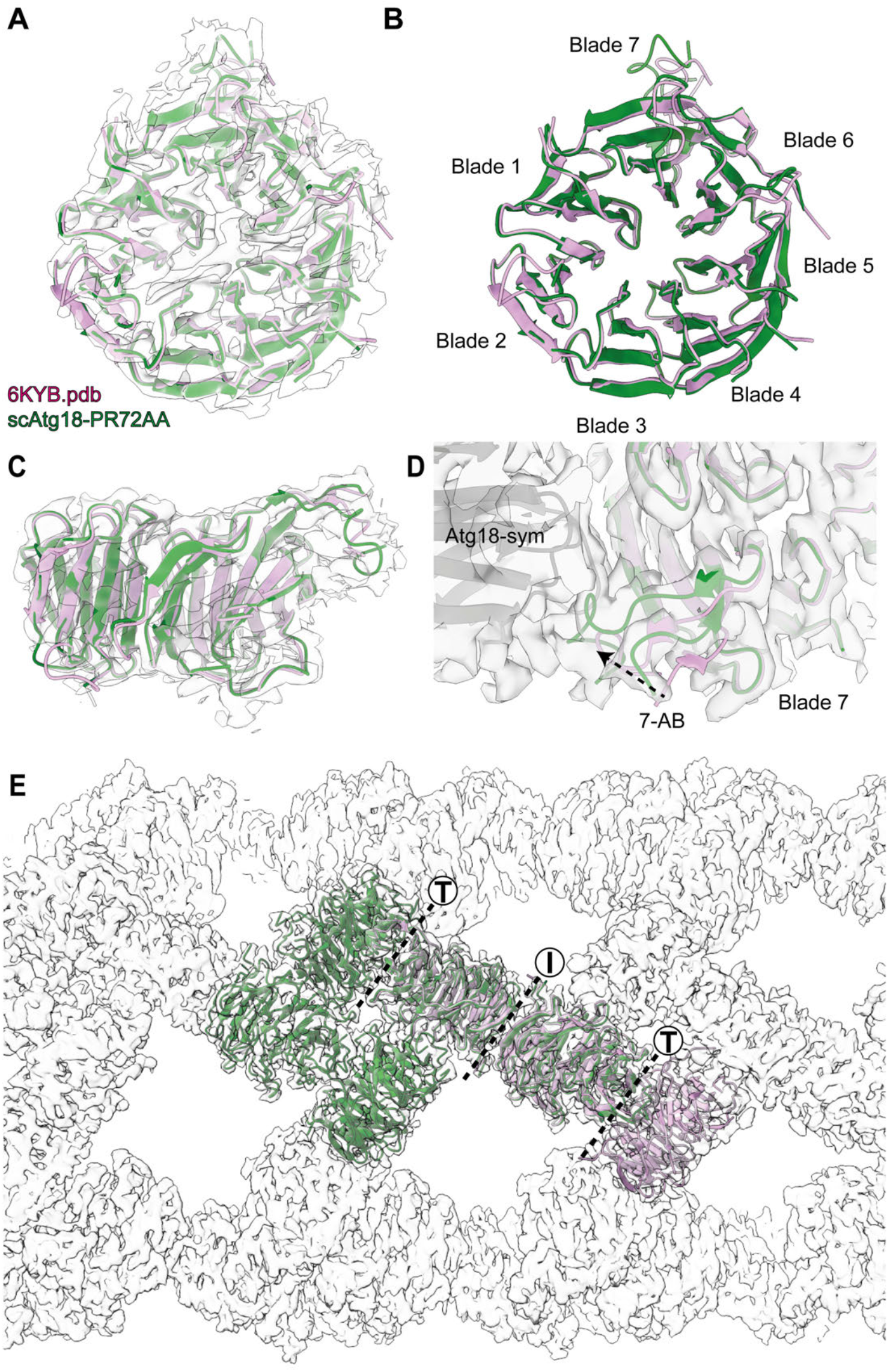
Comparison of Atg18-PR72AA cryo-EM tube structure with X-ray structure of Atg18 (PDB-ID 6KYB). (A) Overall fold is very similar between Atg18-PR72AA (green) and Atg18 (PDB-ID 6KYB) (magenta) that matches the experimental cryo-EM density (grey). Superposition in (B) top and (C) side view shows minor differences in the 7AB loop in blade 7. (D) Close up reveals that loop 7AB is bound more closely to the symmetry neighbor in the tubular assembly (grey). (E) 6KYB PDB structure (magenta) contains the same I and T-interfaces as the determined tubular cryo-EM structure (green). Cryo-EM density of the helical tube is displayed in grey. 6KYB PDB structure was structurally aligned to the Atg18-PR72AA atomic model using the matchmaker program in ChimeraX.

**Suppl. Fig. 3.**
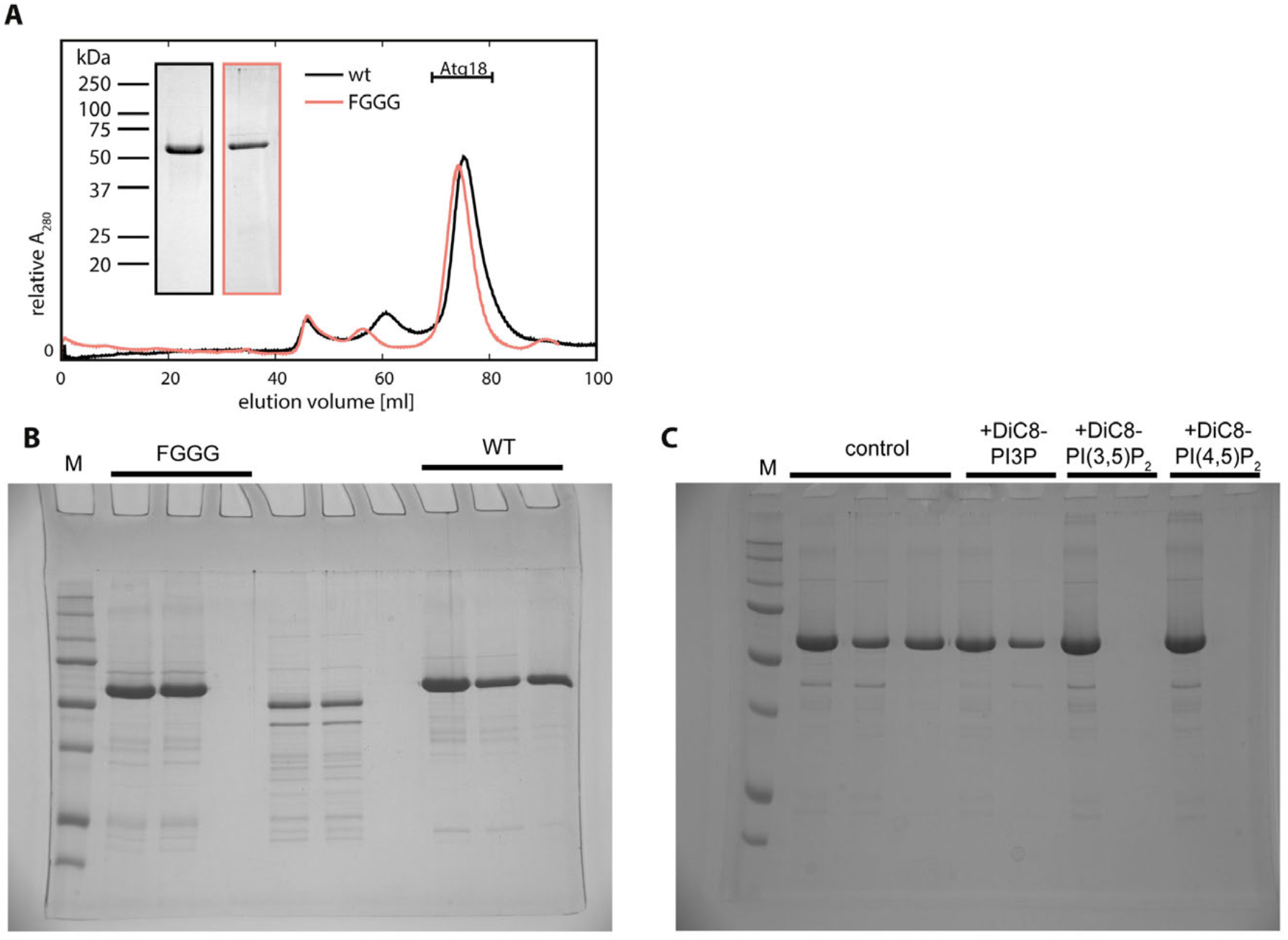
Gel filtration profiles and pelletation assay of Atg18-WT and Atg18-FGGG. (A) Gel filtration profile of Atg18-WT (black) and Atg18-FGGG (rose). Uncropped and unedited Coomassie-stained SDS-PAGE gels of pelletation assay used for preparing Fig. 3: (B) FGGG mutant (left) in comparison with Atg18-WT (left) and (C) Atg18 incubated with water (control), soluble DiC8-PI3P, DiC8-PI(3,5)P_2_ and DiC8-PI(4,5)P_2_ as indicated on top of the lanes.

**Suppl. Fig. 4.**
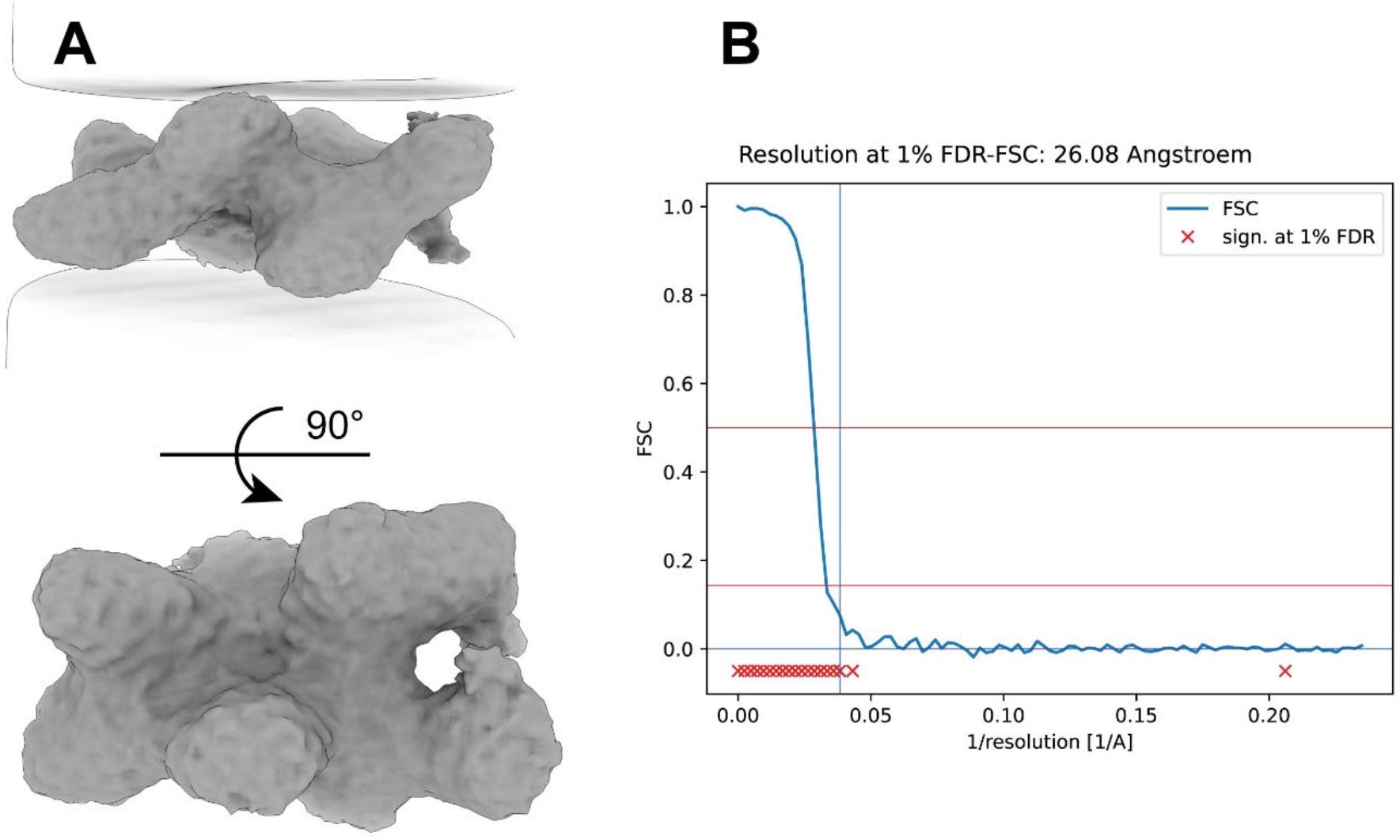
Subtomogram average structure of membrane-bound Atg18. (A) Subtomogram average density of membrane-bound Atg18 and (B) the corresponding resolution determination by FSC at 1% FDR.

**Suppl. Fig. 5.**
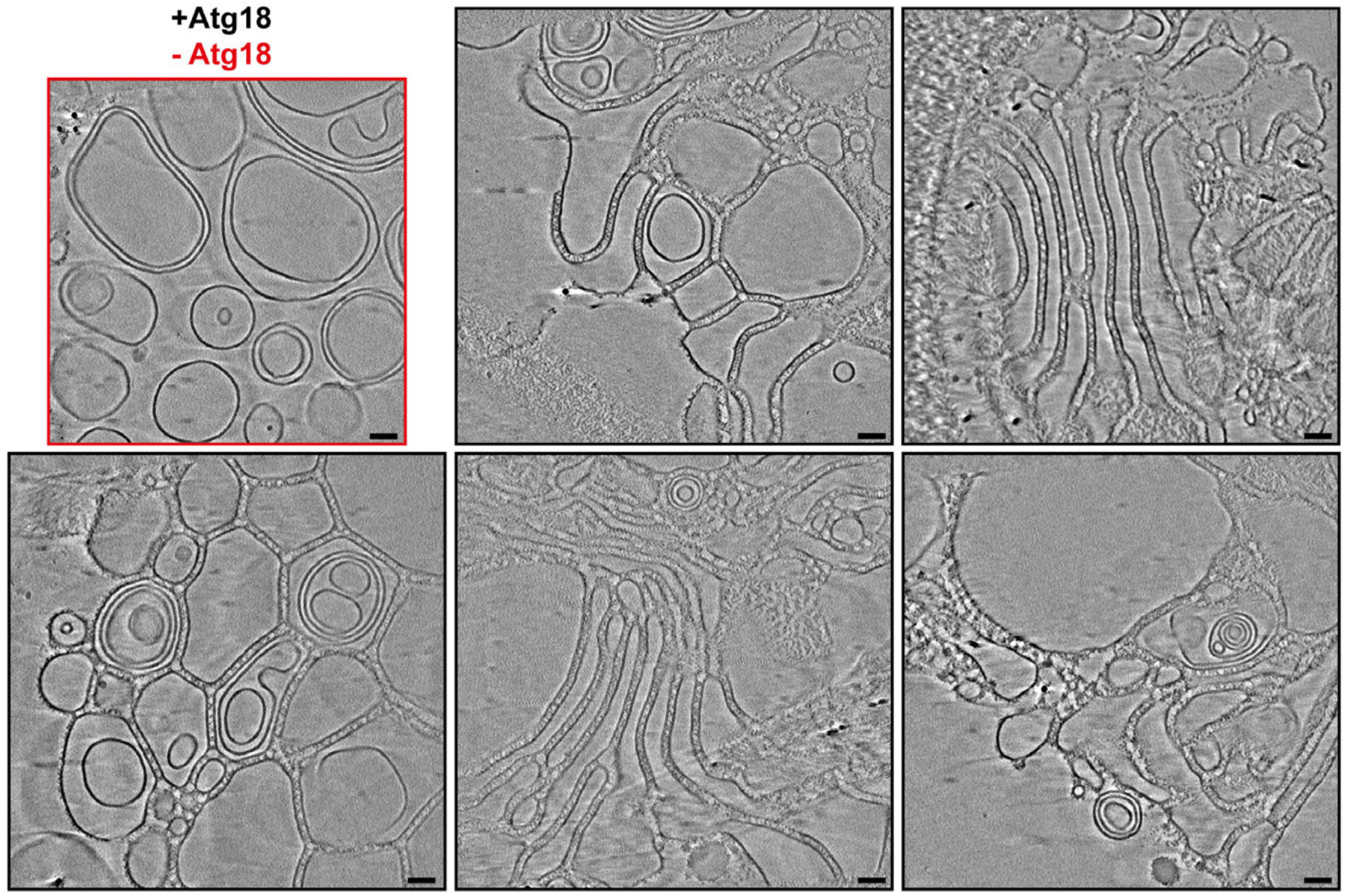
Tomogram slices of Atg18 with PI(3,5)P_2_-containing large unilamellar vesicles (LUVs). Reconstructed tomogram slices of LUVs alone (red box, top left) and with incubated Atg18 (black boxes): different examples of observed lipid morphologies that share long stretches of juxtaposed parallel lipid membranes with inter-membrane connecting densities. Scale bar corresponds to 50 nm.

**Suppl. Fig. 6.**
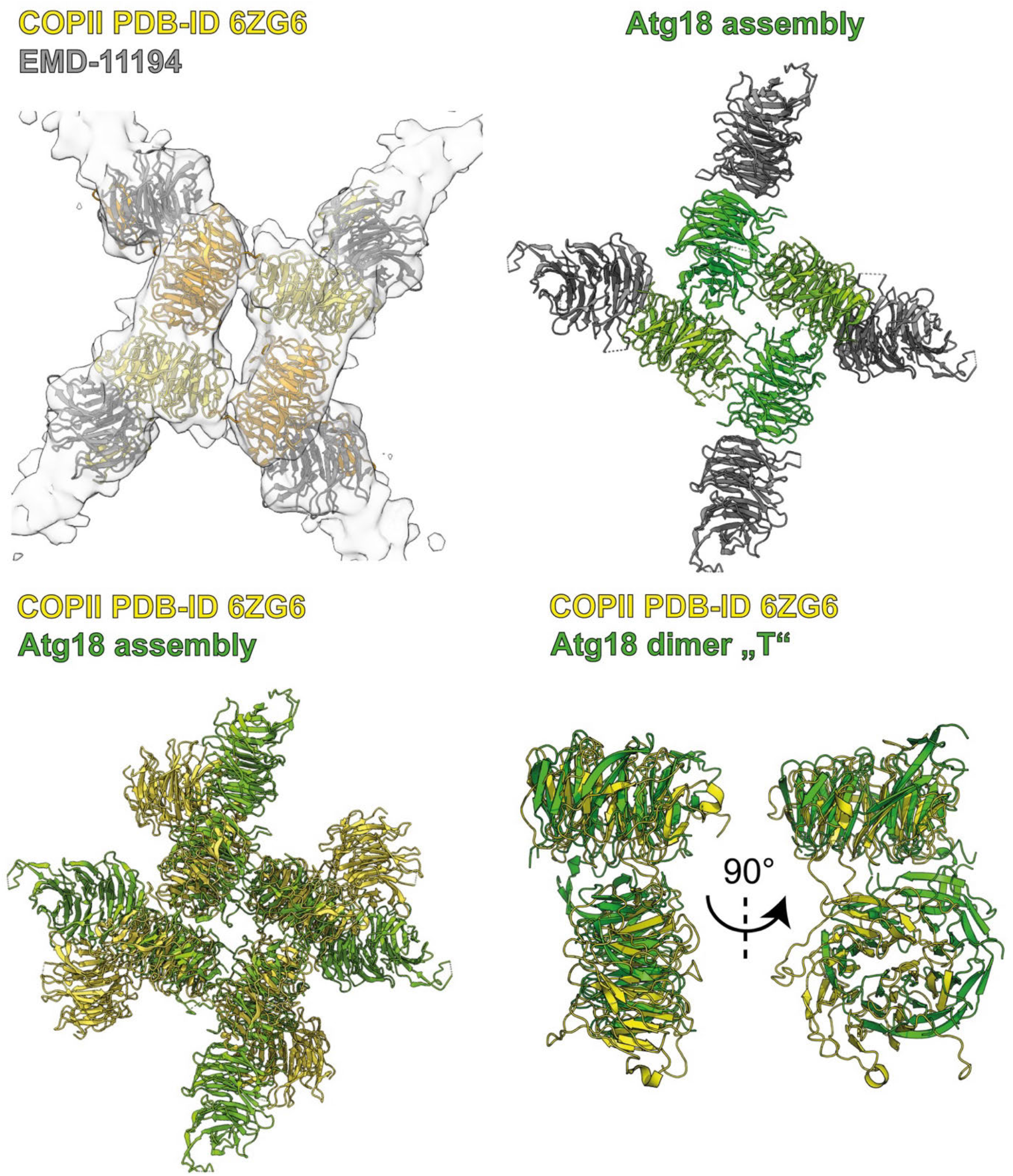
Structural comparison of outer coat COPII with Atg18 tubular assembly. Top. Side views of lozenge pattern for COPII/Sec13-Sec31 (PDB ID 6ZG6) with corresponding cryo-EM density (EMD-11194) (left, yellow) and Atg18 (right, green). Bottom left. Side view of superimposed diamond-shaped COPII/Sec13-Sec31 (yellow) and Atg18 (green) atomic models. Bottom right. T-interface superposition of atomic models of COPII/Sec13-Sec31 (yellow) and Atg18 (green).

**Supp. Movie 1: reconstructed and denoised tomogram from Fig. 5**

https://erc-3-cloud.fz-juelich.de/s/7Tye4AQY4dAa5Fx

